# WHIRLY1 regulates aliphatic glucosinolate biosynthesis in early seedling development of Arabidopsis

**DOI:** 10.1101/2024.04.12.589156

**Authors:** Linh Thuy Nguyen, Pinelopi Moutesidi, Jörg Ziegler, Anike Glasneck, Solmaz Khosravi, Steffen Abel, Götz Hensel, Karin Krupinska, Klaus Humbeck

## Abstract

WHIRLY1 belongs to a family of plant-specific transcription factors capable of binding DNA or RNA in all three plant cell compartments that contain genetic materials. In *Arabidopsis thaliana*, WHIRLY1 has been studied at the later stages of plant development, including flowering and leaf senescence, as well as in biotic and abiotic stress responses. In this study, *WHIRLY1* knock-out mutants of *A. thaliana* were prepared by CRISPR/Cas9 to investigate the role of AtWHIRLY1 during early seedling development. The loss-of-function of *WHIRLY1* in 5-day-old seedlings did not cause differences in the phenotype and the photosynthetic performance of the emerging cotyledons compared to the wild type. Nevertheless, comparative RNA sequencing analysis revealed that the knock-out of *WHIRLY1* affected the expression of a small but specific set of genes during this critical phase of development. About 110 genes were found to be significantly deregulated in the knockout mutant, wherein several genes involved in the early steps of aliphatic glucosinolate (aGSL) biosynthesis were suppressed compared to wild type plants. The downregulation of these genes in *WHIRLY1* knock-out line led to a decreased GSL contents in seedlings and in seeds. We also examined myrosinase activity during seed-seedling transition and showed that the reduction in aGSL biosynthesis is the main reason for lowering aGSL content in young seedlings. The results suggest that AtWHIRLY1 plays a role in regulating aliphatic glucosinolate biosynthesis during early seedling development.

**Significance statement:** WHIRLY1 functions in several aspects of plant development and stress responses, however little is known about its involvement in young seedling development. Here we show that in this stage, WHIRLY1 specifically regulates expression of genes encoding enzymes in the early steps of aliphatic glucosinolate biosynthesis pathway, leading to a reduction in glucosinolate content in the *WHIRLY1* knock-out seedlings.

## Introduction

The WHIRLY protein family comprises few plant-specific proteins sharing the eponymous single-stranded (ss)-DNA-binding domain Whirly (Desveaux et al., 2000; Desveaux et al., 2002). The members of this family have the ability to bind preferentially to ssDNA but can also bind to dsDNA and RNA (Desveaux *et al*., 2002; Cappadocia *et al*., 2010). Each WHIRLY protein contains an N-terminal organelle target peptide (OTP), which targets them into different cellular compartments. WHIRLY1 was reported to be targeted to plastids (Isemer et al., 2012 a; Desveaux et al., 2005) and also found in nucleus (Grabowski *et al*., 2008). By its dual localization in chloroplasts and nucleus, WHIRLY1 is an excellent candidate for the communication between organelles and the nucleus during plant development and stress responses (Krupinska *et al*., 2022; Taylor *et al*., 2022).

The first discovered WHIRLY protein was potato WHIRLY1, which was shown to form a whirligig-like homotetramer called PBF-2. The binding of PBF-2 to the elicitor response element (ERE; TGACAnnnnTGTCA) in the promoter of the potato pathogenesis-related gene 10a (*StPR-10a*) induced its expression in response to pathogen attack (Desveaux et al., 2000; Desveaux et al., 2002). Similarly, Arabidopsis WHIRLY1 can bind to the promoter sequence of *PR1* and thereby influence plants’ defense (Desveaux *et al*., 2004). In Arabidopsis *WHIRLY1* TILLING mutants, mutated WHIRLY1 versions showed attenuated binding ability toward *PR1* promoter, coinciding with reduced resistance towards *P. parasitica* (Desveaux *et al*., 2004). Moreover, WHIRLY1 functions in response to biotic stresses were observed in other species, such as maize (Kretschmer *et al*., 2017), cassava (Liu *et al*., 2018), etc.

WHIRLY1 can bind to an ERE-like element overlapping with a W-box in the promoter of *HvS40* in barley (Krupinska *et al*., 2014) and also to the GNNNAAATT sequence plus an AT-rich telomeric repeat-like sequence in *AtWRKY53* promoter in *A. thaliana* (Miao *et al*., 2013). Both genes function in leaf senescence (Humbeck *et al*., 1996; Miao *et al*., 2004) and are proposed to be negatively regulated by WHIRLY1. In addition to functions in pathogen response and leaf senescence, WHIRLY proteins are involved in abiotic stress responses, e.g., drought, possibly via modulating histone markers in the promoter of stress-related genes including *HvNCED1* (Janack et al., 2016; Manh et al., 2023). In fact, all three WHIRLIES in Arabidopsis were found to be associated to epigenetic reprogramming of meristem tissues in root and shoot (McCoy *et al*., 2021). Furthermore, AtWHIRLY1 was shown to interact with HDA15 to modify histone markers of several flowering-related genes (Huang *et al*., 2022).

Function of WHIRLY1 in early development was studied in monocots. *WHIRLY1* mutants of maize and rice exhibited severe disturbances in chloroplast development and consequently died at the three-to-four-leaf stage (Prikryl *et al*., 2008; Qiu *et al*., 2022). In contrast, neither T-DNA insertion mutants of Arabidopsis *WHIRLY1* nor *WHIRLY2* show apparent phenotypes. It has been, therefore, proposed that AtWHIRLY3 might replace AtWHIRLY1 and AtWHIRLY2 (Krupinska *et al*., 2022). Analysis of the *why2* mutant showed that germination and early seedling development were compromised (Golin *et al*., 2020; Negroni *et al*., 2024), whereas vegetative growth was unaffected. This result indicates that AtWHIRLY3 cannot replace all functions of AtWHIRLY2 during germination, which is in line with the low transcript abundance of *AtWHIRLY3* at this stage of development (Golin *et al*., 2020).

So far, research on the biological functions of Arabidopsis WHIRLY1 mainly concerned senescence and flowering. Still, little is known about the involvement of WHIRLY1 in early seedling development, which plays a vital role in the growth cycle. Therefore, this study aims to investigate whether AtWHIRLY1, similarly to AtWHIRLY2, plays a role in the early stage of plant’s life. Most previous studies on the function of AtWHIRLY1 used T-DNA insertion mutant *why1-1* (SALK_023713) featuring a T-DNA integrated in a close proximity to the start codon (Isemer et al., 2012 b; Huang et al., 2022; Lin et al., 2019). Whereas, in the present work, new *AtWHIRLY1* knock-out mutant lines prepared by site-directed mutagenesis CRISPR/Cas9 approach were used to study functions of WHIRLY1 at the early stage of development. The loss-of-function of *AtWHIRLY1* did not cause apparent changes regarding phenotype and photosynthetic performance of young seedlings. However, genome-wide gene expression analysis revealed a significant impact of *WHIRLY1* loss on the expression of a small set of nuclear genes. These genes encode enzymes involved in early steps of aliphatic glucosinolate (GSL) biosynthesis, leading to a lower aliphatic GSL abundance in the *AtWHIRLY1* knock-out mutant compared to wild type seedlings.

## Results

### CRISPR/Cas9-mediated knock-out mutants of *AtWHIRLY1*

So far, most studies about the AtWHIRLY1 function used the mutant *why1-1* with a T-DNA insertion in the first exon (SALK_023713), which had been first denominated KO-1 (Yoo *et al*., 2007). However, in this mutant and two other T-DNA insertion mutants, i.e., *why1-2* (SALK_147680; KO-2 in Yoo et al., 2007 and *why1-3* (SALK_03900), mRNA level of *AtWHIRLY1* was relatively high (Figure S1, Schaller, 2017). Noticeably, the *why1-1* mutant still has a second in-frame ATG codon located downstream of the first ATG which might serve as an alternative start codon for the translation of a shorter protein lacking the first 10 amino acids of the plastid transit peptide. Hence, this truncated form of WHIRLY1 might be synthesized in the *why1-1* mutant but mistargeted. To preclude this possibility, *AtWHIRLY1* knock-out mutants were prepared by CRISPR/Cas9-mediated genome editing in the ecotype Col-0 background. Three independent Cas9-free homozygous mutant lines were generated and denotated as *why1-4*, *why1-5*, and *why1-6*. *why1-4 and why1-5* mutants contain point mutations, 25_27delAA and 27_28insT, respectively, shifting the reading frame and altering the amino acid sequences of the residual proteins having 50 instead of 263 amino acids (Figure 1a). Whereas, the mutation in the line *why1-*6 resulted in a novel in-frame ATG codon after the 27_28insA mutation which might produce a truncated WHIRLY1 protein. To avoid the effects of such an isoform, this mutant was not investigated in this study. The two mutant lines, *why1-4* and *why1-5*, were verified using PCR-coupled cleavage amplified polymorphism site analysis (Figure 1b), showing that they are homologous to the mutated alleles. Besides, the absence of a Cas9-coding sequence was checked by PCR with specific primers (Table S4), showing that both lines are Cas9-free. The knock-out of *WHIRLY1* by CRISPR/Cas9 resulted in a substantial reduction of *WHIRLY1* transcript level compared to the wild type (Figure 1c), whereas expression levels of the two others *WHIRLY* genes were similar as in the wild type plants.

**Figure 1.**
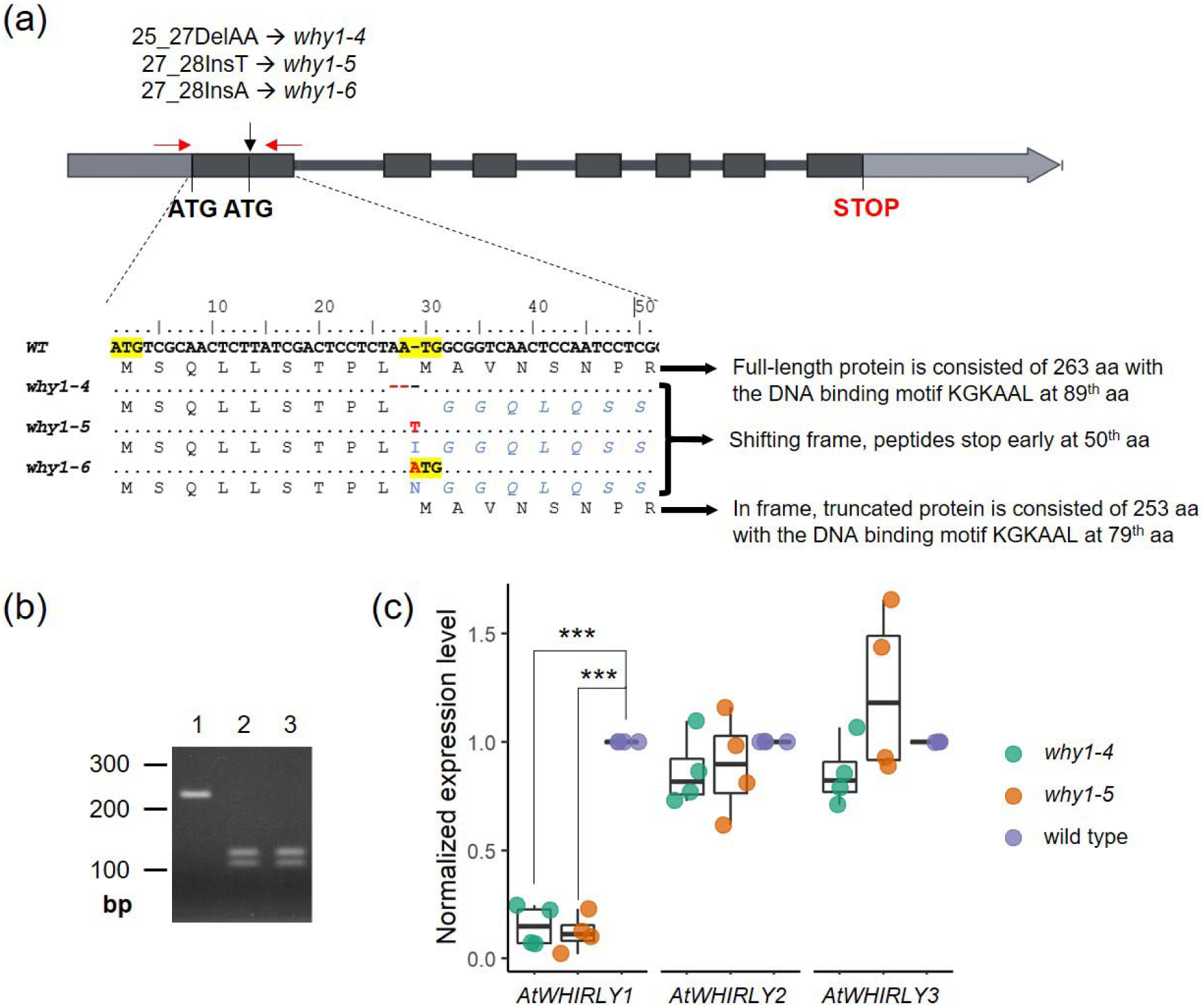
Genotyping of CRISPR/Cas9 mutants of *AtWHIRLY1*. (a) Scheme of three WHIRLY1 knock-out mutants in the present work. The start codon (ATG) is highlighted and point mutations are depicted in red letters. Both *why1-4* and *why1-5* lines contain frame-shift mutations, which change the protein sequence starting at 10^th^ aa. The mutation in w*hy1-6* might also cause frame shifting together with a novel ATG in frame within the wild type allele. Sizes of the expected mutant peptides encoded by these alleles are shown at the end of the alignment. (b) Two mutants, *why1-4* and *why1-5,* were verified using PCR-coupled CAPS. The wild-type allele gave a non-cut fragment of 231 bp while CRISPR/Cas9-mediated mutated alleles were digested into two products of 126 bp and 105 bp. The homologous mutants produce only two smaller bands. Lane 1: wildtype, lane 2: *why1-4*, lane 3: *why1-5*. (c) Normalized expression level of three *WHIRLY* genes in three knock-out mutants by qRT-PCR compared to wild type plants. Boxplot central line shows median value, box limits indicate the 25^th^ and 75^th^ percentile, excluding outlier(s). Whiskers extend 1.5 times the interquartile range. Each dot represents one individual datapoint as one of four independent biological replicates, each containing about 30 seedlings. Asterisks denote a statistically significant level of Student’s t-test between qRT-PCR-based normalized expression level of genes in *WHIRLY1* mutants and the wild type plants, (***) p-value < 0.001.

### Loss-of-function of *WHIRLY1* did not affect morphology and photosystem II efficiency

To investigate the impact of *WHIRLY1* loss on seedling development, comparative phenotypic analysis of CRISPR/Cas9 knock-out lines (*why1-4* and *why1-5*) and the wild type was performed. After five days grown on half-strength MS medium, seedlings developed roots and green cotyledons (Figure 2a). Morphologically, there was no apparent difference between all lines regarding root length, hypocotyl length, and fresh weight (Figure 2b). Photosystem II efficiency indicated by chlorophyll fluorescence (F_V_/F_M_) and chlorophyll content of cotyledons were similar between wild type and mutant seedlings (Figure 2b). The results indicate that during early development under normal growth conditions, the loss-of-function of *WHIRLY1* has no evident influence on the vegetative growth and development of photosynthetic capacity.

**Figure 2.**
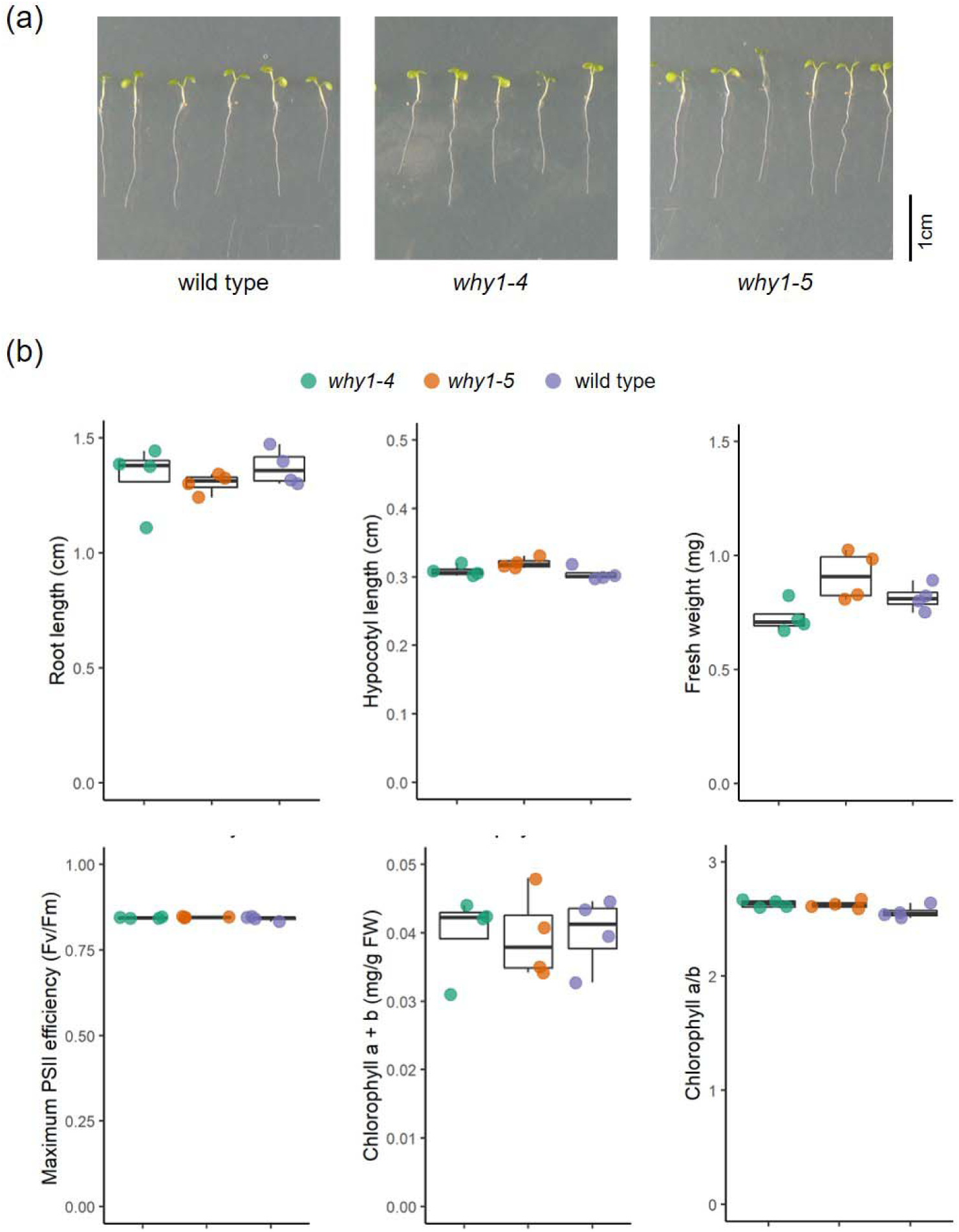
Phenotype of *WHIRLY1* knock-out mutant seedlings. (a) Image of 5-day-old wild type and two CRISPR/Cas9 knock-out mutants of *AtWHIRLY1*, *why1-4* and *why1-5*. (b) Root length, hypocotyl length, whole seedling fresh weight, maximum PSII efficiency (Fv/Fm), total chlorophyll a+b content, and chlorophyll a/b ratio of 5-day-old WT and *why1* seedlings. Boxplot central line shows median value, box limits indicate the 25^th^ and 75^th^ percentile, excluding outlier(s). Whiskers extend 1.5 times the interquartile range. Each dot represents one individual datapoint as one of four independent biological replicates, each containing about 20-30 seedlings. Asterisks indicate statistically significant level based on the Student’s t-test, *** p-value < 0.001.

### Loss-of-function of *WHIRLY1* specifically influenced the transcriptome of seedlings

To examine how absence of functional *AtWHIRLY1* influences nuclear gene expression in 5-day-old seedlings, comparative transcriptome analysis by RNA sequencing (RNA-seq) was performed between the *WHIRLY1* knock-out line *why1-5* and the wild type plants. Total RNA isolated from around 30 seedlings grown on the same agar plate served as one biological replicate. The approach consists of three independent biological replicates per sample. A summary of read numbers and read mapping rate is presented in Table S1.

In each line, nearly 15 thousand genes were expressed (with an expression level above the threshold, i.e., fpkm value > 1). The differences in gene expression between *why1-5* and wild type seedlings were identified using DESeq2 program. Considering that mutant seedlings did not show apparent changes in morphology and photosystem II efficiency compared to the wild type plants, differentially expressed genes (DEGs) were evaluated using a mild stringent cut-off of |log_2_fold-change(FC)| > 1 and p-value < 0.01. As a result, 73 upregulated DEGs and 42 downregulated DEGs were identified in the *why1-5* mutant compared to wild type seedlings, indicating that WHIRLY1 can regulate target genes both positively and negatively. The low number of DEGs (Figure 3a) also implies that the loss-of-function of *WHIRLY1* has no profound influence on the transcriptome during this early growth stage, but rather has a specific impact on a small gene set. Gene Ontology (GO) enrichment analysis failed to identify significant GO terms for over-accumulated genes in *why1-5* compared to wild type seedlings. On the other hand, a small set of 42 significantly downregulated DEGs due to *WHIRLY1* knock-out was enriched with genes related to the biosynthesis of sulfur-containing compounds. In particular, the glucosinolate (GSL) biosynthesis ontology term showed a rather high enrichment fold (Figure 3b). Besides GO enrichment analysis, a manual search for all upregulated and downregulated DEGs in public databases (The Arabidopsis Information Resource (TAIR), www.arabidopsis.org and UniProt, https://www.uniprot.org/) revealed that multiple DEGs due to *WHIRLY1* loss are associated to various other processes, including development and stress responses (Table S2).

**Figure 3.**
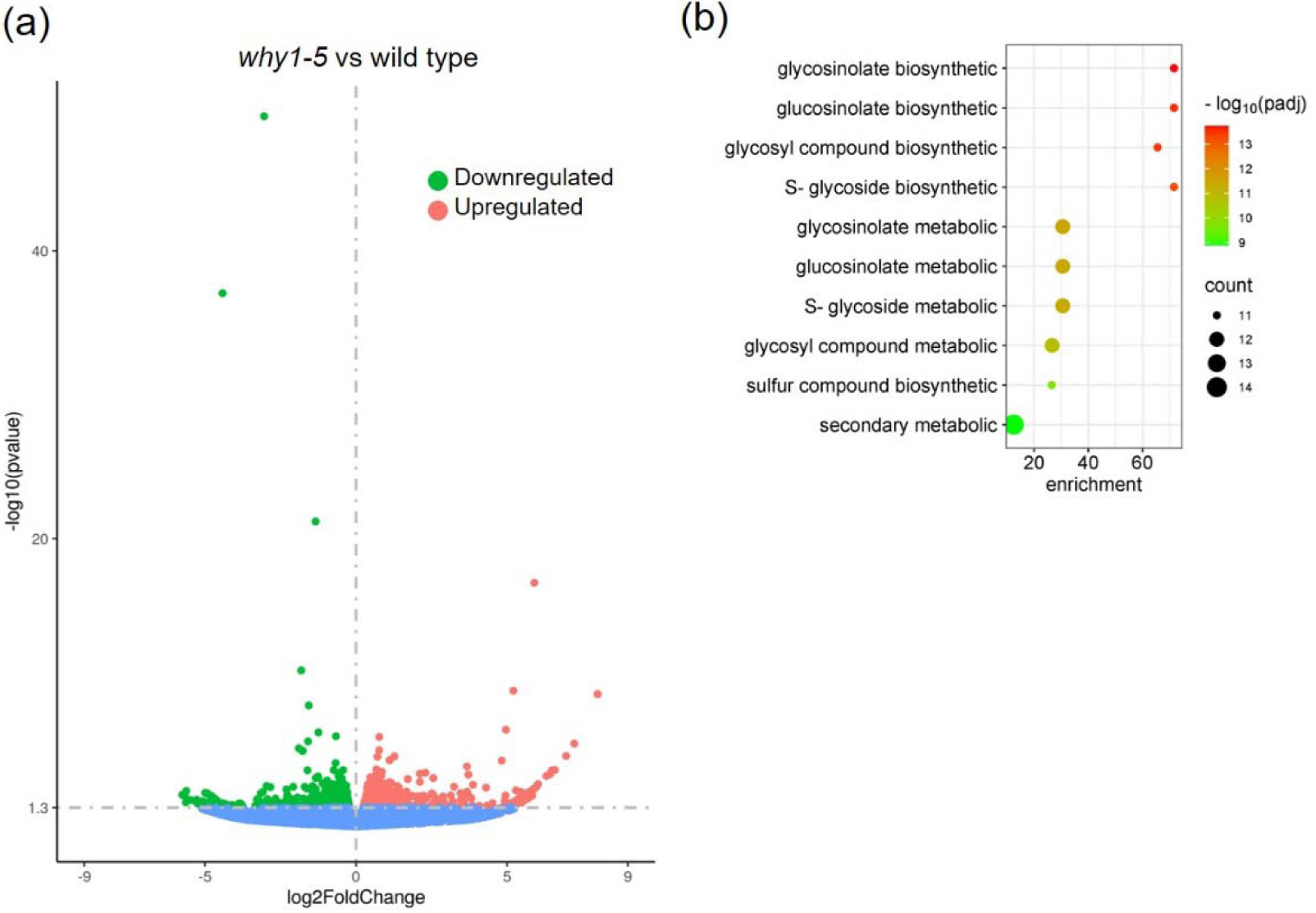
Comparative transcriptomic analysis between *why1-5* mutant and wild type seedlings. (a) The expression changes of DEGs in the *why1-5* mutant compared to the wild-type seedlings are shown by a Volcano plot. (b) GO enrichment analysis of downregulated DEGs. The top 10 GO biological process terms according to p-adjusted value are shown.

### Downregulation of genes encoding enzymes of the aliphatic glucosinolate pathway

A striking result of the comparative transcriptomic analysis was that 10 downregulated DEGs are involved in glucosinolate (GSL) biosynthesis (Figure 3b). GSLs are secondary metabolites which are composed of a thioglucose, a sulfate group, and a variable side chain derived from amino acids (Mérillon et al., 2017). With regard to the precursor amino acid, GSLs are categorized into three groups: aliphatic glucosinolate (aGSL), indole glucosinolates (iGSL), and benzenic glucosinolate (bGSL). As illustrated in Figure 4a, aGSL biosynthesis involves three main stages: side-chain elongation occurring in the plastid, core-structure formation and side-chain modification which both taking place in the cytosol.

**Figure 4.**
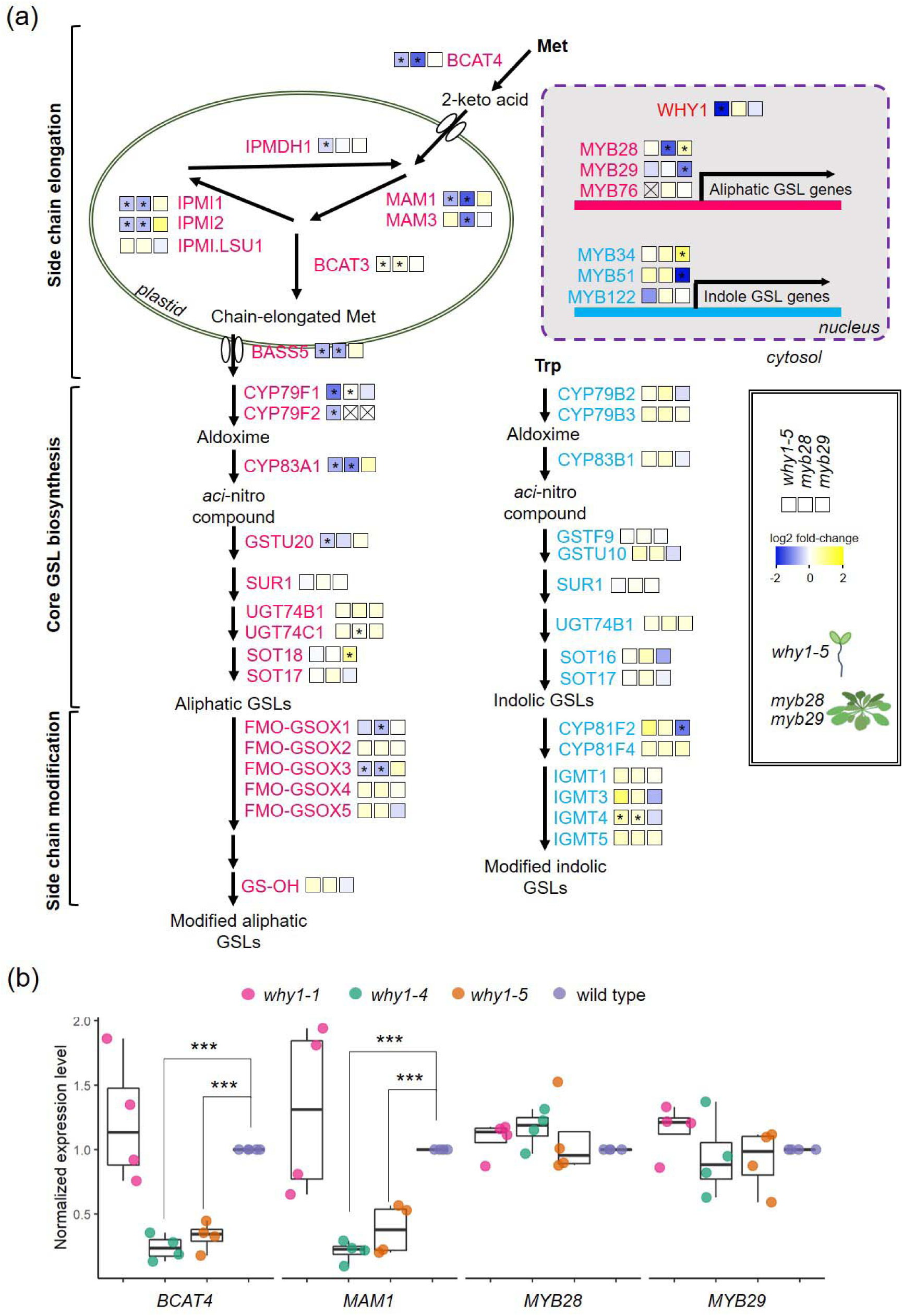
Loss-of-function of *WHIRLY1* affects GSL-related gene expression. (a) Heatmap showing changes in transcript levels of GSL-related genes due to the knock-out of *WHIRLY1* and due to knock-out of *MYB28* and *MYB29* (Ida Elken Sønderby et al., 2010). The log_2_FC is indicated by color bar, and asterisks (*) show statistically significant differences as a result of the RNA-seq or microarray. The scheme of two major GSL biosynthesis pathways, aliphatic GSL and indole GSL, and also for gene names have been adapted from Augustine & Bisht, 2017. See there also for gene abbreviations. (b) Normalized expression levels of selected GSL-related genes measured by qRT-PCR. Boxplot central line shows median value, box limits indicate the 25^th^ and 75^th^ percentile, excluding outlier(s). Whiskers extend 1.5 times the interquartile range. Each dot represents one individual datapoint as one of four independent biological replicates, each containing about 30 seedlings. Asterisks indicate statistically significant level of Student’s t-test between *WHIRLY1* knock-out mutants and WT; * p-value < 0.05, ** p-value < 0.01, *** pvalue < 0.001.

According to the transcriptomic data, loss-of-function of *WHIRLY1* specifically affected the transcript abundances of several genes encoding enzymes in the early steps of Methionine (Met)-derived aGSL biosynthesis, such as *BRANCHED-CHAIN AMINOTRANSFERASE 4* (*BCAT4*), *METHYLTHIOALKYLMALATE SYNTHASE 1* (*MAM1*), *ISOPROPYLMALATE ISOMERASE 1* (*IPMI1*), and *BILE ACID:SODIUM SYMPORTER 5* (*BASS5*), wherein most of them are plastid-localized (Figure 4a). Additionally, genes encoding enzymes in later stages of GSL biosynthesis were also suppressed in the *why1-5* mutant compared to the wild type seedlings, such as *CYTOCHROME P450 79F1 (CYP79F1)*, *CYP79F2,* and *CYP83A1* (Figure 4a). The downregulation of the aforementioned genes due to *WHIRLY1* knock-out was validated by qRT-PCR, showing a high correlation between RNA-seq and qRT-PCR data (Figure S2). The expression levels of numerous genes related to biosynthesis of aGSL in the *why1-5* seedlings were substantially decreased to about a third of the wild-type levels (Figure S2). In contrast, the expression of most genes involved in iGSL biosynthesis was not affected in the *WHIRLY1* knock-out mutant compared to wild type plants (Figure S2). Still, *SUPERROOT1* (*SUR1*), a gene shared between aGSL and iGSL pathways, showed slightly reduced transcript levels in the *why1-*5 compared to wild type seedlings. The downregulation of selected aGSL biosynthetic genes was also confirmed in seedlings of the other CRISPR/Cas9-mediated knock-out line *why1-4* (Figure 4b). Interestingly, in contrast to the CRISPR/Cas9 knock-out mutants, expression of GSL-related genes in the T-DNA insertion mutant *why1-1* was unaltered compared to wild type plants (Figure 4b).

Biosynthesis of aGSL is known to be directly controlled by MYELOBLASTOSIS transcription factors (MYBs), including MYB28, MYB29, and MYB76, while iGSL biosynthesis is governed by MYB34, MYB51, and MYB122 (Mérillon et al., 2017). Therefore, the expression levels of the corresponding genes were also analyzed in the *why1-5* mutant and wild type seedlings, showing a similar transcript level of *MYB* genes in both lines (Figure 4b; Figure S2). The findings indicate that WHIRLY1 does not regulate aGSL biosynthesis genes by indirectly modulating transcription of these upstream regulators, i.e., *MYB28* and *MYB29*, but likely via another mechanism.

Transcription alteration of GSL-related genes due to *WHIRLY1* loss in seedlings was then compared to the expression changes of corresponding genes because of *MYB28* and *MYB29* knock-out in mature plants using the available RNA-array data (Sønderby et al., 2010). Interestingly, similar changes in GSL biosynthesis gene expressions were observed in the *myb28* and *why1-5* line compared to wild type plants (Figure 4a). In contrast to these two mutants, the *myb29* showed a different pattern in the expression of GSL-related genes. Indeed, the same genes involved in side-chain elongation steps were repressed due to the loss-of-function of either *WHIRLY1* or *MYB28*, but not *MYB29*. For instance, *BCAT4*, which encodes the key enzyme initiating the side-chain elongation process, was significantly suppressed in the knock-out lines *why1-5* and *myb28* compared to wild type plants. Meanwhile, *BCAT4* transcript levels did not alter due to *MYB29* knock-out. On the contrary, genes encoding enzymes in the iGSL pathway were slightly but not significantly induced in both *why1-5* and *myb28* lines compared to the wild type, while some genes, such as *MYB34* and *CYP81F2*, were significantly deregulated in *myb29* mutant. Interestingly, the effect of the loss-of-function of WHIRLY1 on aGSL-related gene expressions was only observed in the seedling stage but not in the rosette of mature plants (Figure S3).

### Disturbance in GSL content in the *WHIRLY1* knock-out mutant

To investigate whether the specific suppression of genes encoding enzymes in the early steps of aGSL biosynthesis has an influence on aGSL contents, 5-day-old *why1-5* and wild type seedlings were collected to quantify GSLs by LC-MS/MS. The amount of each GSL in each sample was calibrated using sinigrin, a compound not naturally found in *A. thaliana*, then normalized to sample fresh weight. GSL profiles of the *WHIRLY1* knock-out *why1-5* and wild type seedlings were in line with the transcriptomics data (Figure 5a; Supplement Table S3). In 5-day-old *why1-5* seedlings, the total GSL content was significantly lower than that in wild type plants, as the results of significantly reduced abundances of individual short-chain aGSL compounds (Figure 5a, Supplement Table S3). For example, the wild type contained almost twice as much 4-methylthio-3-butenyl (4MTB) as *why1-5* mutant seedlings (2.92 nmol/mg FW *versus* 5.02 nmol/mg FW; Supplement Table S3). For comparison, long-chain aGSLs were only reduced mildly due to the knock-out of *WHIRLY1* (Figure 5a). Furthermore, indole GSL and embryo-synthesized GSL concentrations were similar between the two lines.

**Figure 5.**
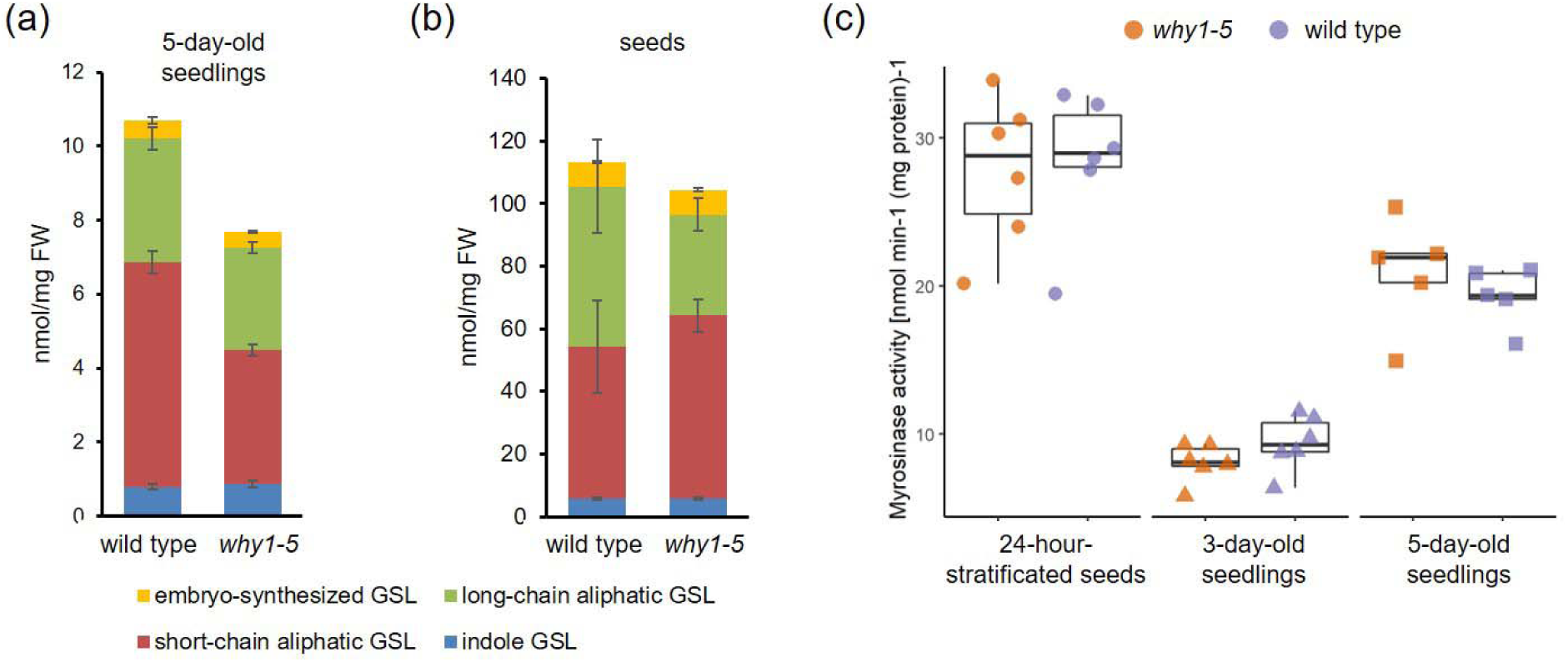
Glucosinolate contents and myrosinase activities in the *WHIRLY1* knock-out and wild type plants. (a, b) Glucosinolate contents (nmol/mg FW) of indole GSL, short-chain aliphatic GSL, long-chain aliphatic GSL, and embryo-synthesized GSL (3-BZOP) measured by HPLC-MS/MS. Bar charts show the average and error bars show standard deviation of (a) three independent biological replicates, each containing four samples as pool of 5-day-old seedlings and (b) six independent samples, each containing around 5 mg seeds. (c) Myrosinase activity [nmol min^-1^ (mg protein)^-1^] of 24-hour-stratificated seeds, 3-day-old, and 5-day-old seedlings. Boxplot central line shows median value, box limits indicate the 25^th^ and 75^th^ percentile, excluding outlier(s). Whiskers extend 1.5 times the interquartile range. Each dot represents one individual datapoint as one of five-six independent biological replicates, each contains around 100 mg of fresh material as a pool of seeds/seedlings.

It has been reported that in young seedlings, majority of GSLs is left over from those synthesized and accumulated during embryo development while minority is newly-synthesized after germination (Meier *et al*., 2019; Jeschke *et al*., 2019). To clarify whether absence of *WHIRLY1* already disturbs GSL composition in seeds, we quantified the GSL contents in *why1-5* and wild type dry seeds (Figure 5b; Table S4). Our data shows that total GSL contents are much higher in seeds than in seedlings (Figure 5b), in line with many other reports (Wittstock and Burow, 2010; Meier *et al*., 2019; Mérillon and Ramawat, 2017). As in seedlings, missing of WHIRLY1 also affected aliphatic GSL in seeds but did not influence indole as well as embryo-synthesized GSLs, leading to an overall lower total GSL concentration in *why1-5* compared to wild type seeds. It suggests that *WHIRLY1* might also act as a positive regulator of aGSL biosynthesis genes in the embryo development stage. Interestingly, short-chain aGSL quantities in the *why1-5* seeds was significantly higher than that in the wild type while long-chain aGSL contents showed an opposite tendency (Figure 5b; Table S4). Perhaps, the conversion to longer chain aGSL is defected in the mutant, resulting in a higher portion of short-chain aGSLs compared to long-chain aGSLs.

### Myrosinase activity during seed-seedling transition is not affected by the lack of *WHIRLY1*

During seed-seedling transition, GSLs stored in seed are turned over to provide building blocks for the early development by the action of myrosinases (Sugiyama and Hirai, 2019; Sugiyama *et al*., 2021; Meier *et al*., 2019). The discrepancy in total GSL content during this transition between *why1-5* and the wild type raised a question whether the lower aGSL concentrations in mutant seedlings is caused by altered biosynthesis in seedlings and/or by differences in myrosinase activity. Therefore, myrosinase activity in 24-hour-stratificated seeds and seedlings 3 and 5 days after germination were measured (Figure 5c). Protein extract from stratificated seeds showed the highest activity of myrosinases. In comparison, seedlings exhibited a lower myrosinase activity compared to seeds, in which the older seedlings seem to have more myrosinases. Interestingly, there was no difference in myrosinase activity between the *why1-5* mutant and the wild type samples. These results indicate that the different aGSL contents between *why1-5* and wild type seedlings are not because of altered GSL degradation but rather due to newly aGSL biosynthesis during seedling development.

### The evolution of WHIRLY in Brassicales order coincides with glucosinolate diversification

The number of WHIRLY genes and proteins has been investigated restricted to a limited number of species (Muti *et al*., 2024; Qiu *et al*., 2022; Oetke *et al*., 2022; Desveaux *et al*., 2002). To date, no phylogenetic analysis of the WHIRLY family has been conducted. Thanks to the increasing number of plant genomes sequenced in recent years, it is feasible to mine WHIRLY sequences and analyze the evolution of the WHIRLY family. Using Whirly domain sequence of AtWHIRLY1 to perform BLAST searches against NCBI and Pfam databases, thousand sequences which share similarity in either peptide sequence or higher structure were identified. After removing duplications and isoforms, and poor-quality sequences, 788 sequences of 309 species were retained. Sequences then were aligned and used to build a phylogeny tree of the WHIRLY family (Figure S4).

The evolutionary tree revealed that in higher plants, WHIRLY proteins can be grouped into two distinctive groups, WHIRLY1-like and WHIRLY2-like (Figure S4). A majority of plant species contain only these two proteins, such as in barley (Grabowski *et al*., 2008), maize (Prikryl *et al*., 2008), and tomato (Akbudak and Filiz, 2019). However, in some lineages, an additional ortholog of either WHIRLY1 or WHIRLY2 has been recorded. It is worth noting that in complex genomes such as polyploidy, number of WHIRLY genes/proteins is higher but generally they do not diverge into more sub-groups (Hu and Shu, 2021). Four lineages of dicots, including Brassicaceae (cabbages), Fabaceae (legumes), Cucurbitaceae (pumpkin), and Salicaceae (poplar), have two distinctive proteins in the WHIRLY1-like group, for instance AtWHIRLY1 and AtWHIRLY3 in *A. thaliana* (Desveaux *et al*., 2002). Meanwhile, only one monocot lineage, Aracaceae (date palm), has two different proteins grouped in a WHIRLY2-like clade (Figure S4). The appearance of additional WHIRLY in such plant groups, that are quite different in physiology characteristics and ecological adaptations, can serve as a source for evolution, such as to gain a new function.

By further investigating the Brassicales order, three WHIRLY proteins are found in numerous genera, including Arabidopsis, Brassica, and Aethionema, which are members of Brassicaceae family, and Tarenaya of Cleomaceae family (Figure 6). Interestingly, in *Carica papaya*, a species of Caricaceae family, only two WHIRLIES are identified (Figure S4). The additional WHIRLY in core families of Brassicales can be explained by the evolutionary history of this plant order. Brassicales were evolutionized thanks to two whole-genome duplication events (WGD), the At-α following the At-β (Edger *et al*., 2018). It is likely that Caricaceae plants do not have additional WHIRLY protein because of its emergence before these WGD events (Figure 6).

**Figure 6.**
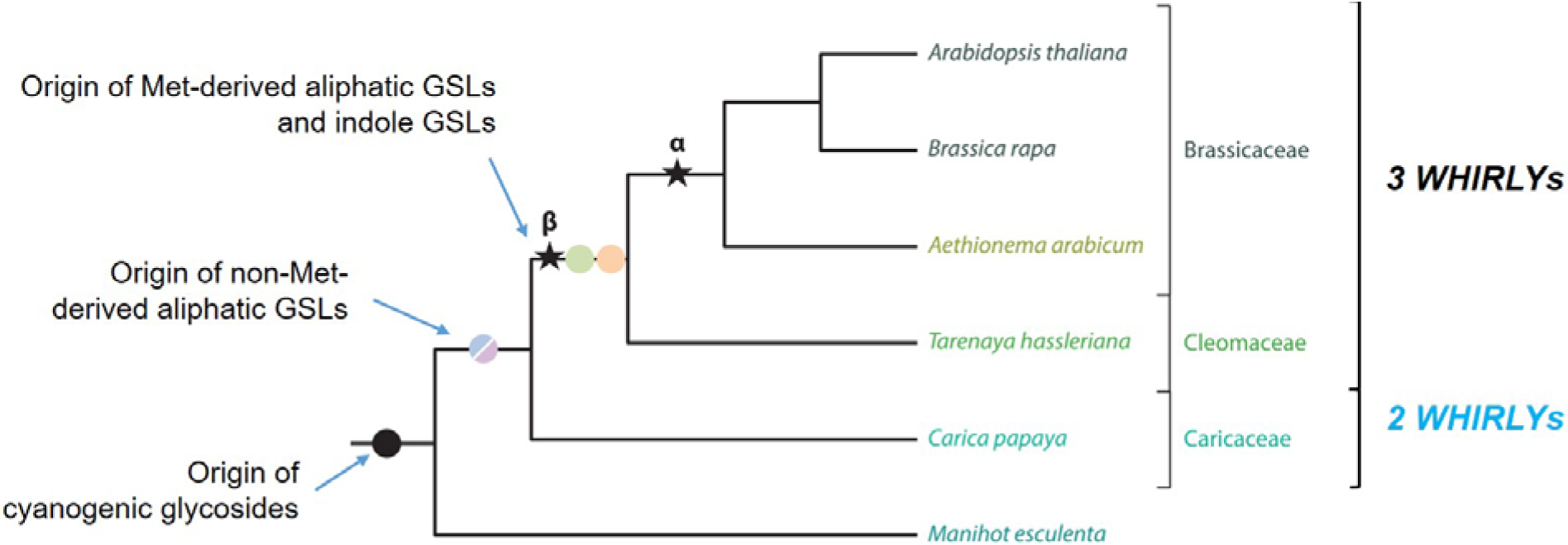
Phylogeny and origin of GSLs. The phylogenetic tree depicted major evolutionary events in GSL diversity uniquely in Brassicales. Circles represent origin of novel GSL groups and stars illustrate WGD events. Number of WHIRLY genes in these families are shown on the right. The scheme was adapted from (Barco and Clay, 2019).

Interestingly, glucosinolates’ diversity in chemical properties is also the results of WGD event and neofunctionalization of genes. Gene duplication at the At-β was the origin of novel iGSL and Met-derived aGSL pathways (Barco and Clay, 2019). Whereas, the recent event At-α gave rise to many side-chain modification genes for both iGSLs and Met-derived aGSLs. Therefore, Met-derived aGSLs only are present in core families of Brassicales, i.e., those derived after the event At-β, such as Brassicaceae, Cleomaceae, etc. (Mérillon et al., 2017). Caricaceae, which has two WHIRLYs, contains only non-Met derived GSL (Mérillon et al., 2017). It is tempting to speculate that the coincidence between additional WHIRLY1-like protein and aliphatic GSL biosynthesis pathway reflects an evolutionary reason for the function of WHIRLY1 in regulation of aGSL biosynthesis.

## Discussion

### Knock-out of *WHIRLY1* has no impact on seedling development but specifically alters gene expressions

WHIRLY1 has been described as a multifaceted transcription regulator that modulates plant development and stress responses (Taylor *et al*., 2022; Krupinska *et al*., 2022). In this study, novel *WHIRLY1* mutants were generated using site-directed mutagenesis with CRISPR/Cas9. These knock-out lines have point mutations which cause a frame shift, changing the protein sequence starting from the 11^th^ amino acid, thereby resulted in relatively short and non-functional peptides (Figure 1a). Despite the true knock-out of *AtWHIRLY1*, even during seedling development, these mutants did not show an obvious phenotype, suggesting that the presence of WHIRLY1 seems unnecessary for average seedling growth. The lack of an apparent phenotype can be explained by an unknown mechanism compensating WHIRLY1-knock-out effects on chloroplast performance and development of seedlings. This complementation in Arabidopsis *WHIRLY1* knock-out mutants could be done preferably by AtWHIRLY3, which was shown to be dually targeted into both chloroplast and mitochondria (Golin *et al*., 2020). It has been observed that the expression level of *AtWHIRLY3* was enhanced in shoots of *why2* knock-out mutant at the 2- to 4-true-leaf stage, suggesting a compensation of WHIRLY2 deficiency by enhanced abundance of WHIRLY3 (Golin *et al*., 2020). With the regard to the dual targeting of WHIRLY3 into chloroplasts and mitochondria, it is likely that AtWHIRLY3 might also replace AtWHIRLY1 in seedling development in *WHIRLY1* knock-out mutants. Indeed, AtWHIRLY1 and AtWHIRLY3 share 77% similarity in protein sequence and might have similar functions under certain conditions (Desveaux *et al*., 2004). For comparison, monocot plants with WHIRLY1 deficiency show obvious phenotypes likely due to lack of a third WHIRLY protein. The RNAi-mediated knockdown *HvWHIRLY1* barley showed delayed formation of green tissue as well as suppressed development of photosynthetic function (Krupinska *et al*., 2019; Saeid Nia *et al*., 2023), while lacking ZmWHIRLY1 in maize or OsWHIRLY1 in rice caused albino seedlings (Prikryl *et al*., 2008; Qiu *et al*., 2022).

Despite the CRISPR/Cas9-mediated *WHIRLY1* knock-out mutants lacking an apparent phenotype, RNA-seq analysis of the *why1-5* mutant seedlings revealed distinct alterations in nuclear gene expression, indicating that the regulation of certain genes is specifically mediated by WHIRLY1. The loss-of-function of *WHIRLY1* affected expression of a small fraction of nuclear genes, which are involved in various developmental processes and stress responses (Table S2). Among differently expressed genes in the *why1-5* mutant compared to the wild type, genes encoding enzymes of aliphatic GSL biosynthesis were significantly suppressed (Figure 4). Interestingly, none of the previously described target genes of WHIRLY1 in mature plants, including *AtPR1* (Desveaux *et al*., 2004) and *AtWRKY53* (Miao *et al*., 2013), were differently expressed in seedlings of the *why1-*5 mutant. This indicates that WHIRLY1 may perform different functions in different developmental stages.

### Participation of WHIRLY1 in regulation of glucosinolate homeostasis during seedling development

Comparative transcriptomic analysis of 5-day-old seedlings revealed that *WHIRLY1* knock-out caused reduced transcript accumulations of several genes involved in aGSL biosynthesis (Figure 4), indicating that WHIRLY1 acts as a positive regulator of these genes. Consequently, aGSL contents were lower in the *why1-5* compared to wild type seedlings (Figure 5a). Majority of seedling GSLs are already synthesized and accumulated in the embryos and serve as a nutrient reservoir for growing seedlings (Wittstock et al. 2010). During germination, GSLs are degraded and sulfur and other nutrients are liberated for the growing seedling (Sugiyama et al. 2021, Meier et al. 2019), leading to substantially reduced GSL concentration during seed-seedling transition. Nevertheless, knock-out of *WHIRLY1* again resulted in a lower GSL accumulation in seeds compared to the wild type (Figure 5b), though the effect of *WHIRLY1* loss in seeds is milder compared to seedlings. Interestingly, *why1-5* seedlings contained reduced amounts of both short-chain and long-chain aGSLs compared to wild type seedlings, whereas *why1-5* seeds showed a significantly higher level of short-chain aGSL but lower level of long-chain compounds than wild type samples (Figure 5a-b). In other words, knock-out of *WHIRLY1* increased the ratio of short-chain to long-chain aGSLs of *why1-5* mutant compared to wild type seeds. This can be explained by a possible attenuation in chain elongation steps of aGSL biosynthesis during embryo development and seed maturation, leading to more short-chain aGSLs being formed than long-chain aGSLs. Given that all GSLs (except embryo-synthesized GSLs) are produced in silique and accumulated into embryos continuously as siliques mature by the action of glucosinolate transporters (Nour-Eldin *et al*., 2012), the measured GSL content in seed reflects both the GSL biosynthesis rate and the transportation capacity. It remains to be investigated whether WHIRLY1 participates in regulation of aGSL biosynthesis and seed GSL accumulation in this developmental stage.

It has been reported that GSL contents in seedlings raise gradually after germination, in which the increase was mainly driven by iGSL followed by long-chain aGSL biosynthesis (Jeschke *et al*., 2019). In another study, GSL quantity increases slightly 2 days after germination (DAG) (in the paper, 4 days on MS medium) followed by a gradual reduction (Meier *et al*., 2019). In another word, shortly after germination, GSL biosynthesis is activated in young seedlings. Missing of *WHIRLY1* clearly reduces the expression levels of genes involved in aGSL biosynthesis during seedling early development (Figure 4), likely lowering newly-synthesized aGSL quantities. Moreover, seed-stored GSLs are hydrolyzed by myrosinases instantly after seed stratification. The protein extract from 24-hour-stratificated w*hy1-5* and wild type seeds showed a quite high myrosinase activity (Figure 5c). After germination, myrosinase activity in young seedlings reduced substantially (Figure 5c), in line with previous report that the GSL content increased from sterile seed to 2 DAG seedling, where likely due to biosynthesis overpass degradation rate (Meier *et al*., 2019). Up to date, preferable GSLs being turned over and their degrading enzymes during seed-seedling transition are still not yet well characterized. Nevertheless, overall myrosinase activities is similar between *why1-5* and wild type samples, suggesting that WHIRLY1 is not involved in regulating GSL hydrolysis via myrosinases. Hence, it is likely that during *WHIRLY1* knock-out seedling development, degradation of seed-stored short-chain aGSLs without re-compensating from *de novo* biosynthesis is the main reason leading to lower these metabolite concentrations in *why1-5* compared to wild type seedlings.

The remaining question is how WHIRLY1 can function in the regulation of the expression of aGSL biosynthesis-related genes in the early development stage. It is known that GSL biosynthesis is regulated precisely at many levels and involves diverse regulators and pathways (Mitreiter et al., 2021). MYB28 is a central transcriptional regulator of aGSL biosynthesis, followed by MYB29 and MYB76 (Sønderby et al., 2010). Interestingly, similar genes were down-regulated due to the knock-out of *WHIRLY1* as well as in the *MYB28* knock-out line (Figure 4a). The influence of knock-out of *WHIRLY1* on GSL metabolism was only observed at the seedling stage, as there was no difference in the expression levels of GSL-associated genes in mature *why1-5* knock-out compared to the wild type plants (Figure S3). This indicates that during early development, WHIRLY1 acts as an additional regulator of aGSL-related genes, while in mature plants, aGSL-related gene expressions are regulated without WHIRLY1 and predominantly by MYB28. It has been reported that WHIRLY1 might bind to melted promoter regions of nuclear target genes, whereby it has an affinity to certain binding motifs, such as the elicitor response element (ERE) including inverted repeat sequences (IR) and W-boxes (Desveaux et al., 2002; Desveaux et al., 2005). Therefore, 1000-bp promoter regions of aGSL genes were analyzed by PLACE (https://www.dna.affrc.go.jp/PLACE/). The analysis revealed different *cis*-elements in the promoters of GSL-related genes. As expected, these promoters contain several MYB binding site motifs for transcriptional regulation by MYB28 and MYB29, and additionally, many W-boxes and IR2 sequences that could serve as potential binding sites of WHIRLY1 are distributed throughout promoter sequences (Figure S5). The binding of WHIRLY1 on promoter of these genes could affect their expression level and/or change epigenetic markers at the region.

### Evolutionary aspects of WHIRLY in combination with glucosinolate diversification

Gene duplication is the primary drive for evolution as it provides material for natural selection. Duplicated genes could either maintain the ancestral function, become silent (pseudogene), or develop a new function. In the present work, the knock-out of *WHIRLY1* strictly affected genes involved in Met-derived aGSL biosynthesis but did not influence the production of other types of GSL. Therefore, the co-occurrence of the novel Met-derived aGSL biosynthesis and an additional *WHIRLY* locus in the genome (Figure 6, Figure S4) prompts the question whether the new WHIRLY protein might have obtained a novel function during evolution. In Brassicaceae, an additional chloroplast-targeted WHIRLY protein might have enabled retrograde control of aGSL biosynthesis as the first steps happen in chloroplast. Synteny analysis showed that *AtWHIRLY1* and *AtWHIRLY3* share the same ancestral gene with monocot *WHIRLY1*, whereby AtWHIRLY3 seems to be more closely related to the monocot WHIRLY1 proteins than AtWHIRLY1 (Krupinska *et al*., 2022). Likely, after the duplication in the event At-β, AtWHIRLY3 kept the original ancestral gene function, while AtWHIRLY1 gained a novel function in the regulation of secondary metabolism. Its evolutionary co-appearance with aGSL biosynthesis-related genes and its dual-localization in chloroplasts and nucleus makes it an ideal candidate regulator of chloroplast-located steps in glucosinolate biosynthesis during seedling development.

## Experimental procedures

### Plant materials and growth

Seeds of *A. thaliana* Columbia-0 and mutants were surface-sterilized in 70% (v/v) ethanol for 5 minutes and then rinsed several times with sterile water. Seeds were placed on Petri plates containing half-strength MS media supplemented with 1% (w/v) sucrose and 0.7% (w/v) agar. After 3 days of stratification in darkness at low temperature, seeds germinated under a continuous light intensity of 15 μE/m^2^s at 22 ± 2°C and 60% relative humidity. Seedlings were investigated at 5 days after germination (DAG). Seedlings were grown vertically and then photographed from the side to determine morphological characteristics. First, a scale was set according to a ruler included in the picture using Fiji ImageJ version 1.53 (Schindelin *et al*., 2012). Hypocotyl and root were marked manually and their lengths were automatically measured.

### Establishment and genotyping of CRISPR/Cas9-mediated knock-out mutants of AtWHIRLY1

Knock-out lines of *AtWHIRLY1* (*At1g14410*) were created by site-directed mutagenesis using CRISPR/Cas9 (Kumlehn *et al*., 2018; Li *et al*., 2021). Annealed oligos were integrated via *Bbs*I restriction enzyme sites in plasmid pSI57 yielding pGH510. The gRNA/Cas9 expression cassettes were introduced into p6i-d35S-TE9 (DNA-Cloning-Service, Hamburg, Germany) via *Sfi*I, generating plasmid pGH482. This construct was used for the *Agrobacterium*-mediated transformation of the ecotype Col-0. Primary transformants (T1) were screened on Murashige and Skoog (MS) media supplemented with 33.3 µg/ml hygromycin B. Resistant seedlings were checked for the presence of DNA encoding the Cas9 endonuclease by PCR with primers Cas9-F2/Cas9-R2. The next generations (T2 and T3) of *Cas9*-positive T1 plants were further analyzed. The mutated *WHIRLY1* locus was screened by PCR amplification and DNA sequencing. The mutated-*WHIRLY1*-containing homozygous lines were also checked for the absence of the Cas9 construct using specific primers.

For genotyping of the *WHIRLY1* knock-out mutants, a PCR-coupled cleavage amplified polymorphism site (CAPS) approach was used. Firstly, segments containing mutated site were amplified by PCR, whereby each reaction comprises 1 µl of diluted DNA (100 ng) as the template, 2 µl 10X *Taq* DNA Polymerase Buffer, 0.4 µl 25 mM MgCl2, 1.2 µl each primer including Why1_F1 and Why1_R1 (Table S4), and 0.1 µl 10 U/µl *Taq* DNA Polymerase (EURx, Poland). Then, 5 µl PCR products were digested with restriction enzyme in a 20 µl reaction with 2 µl 10X NEBuffer 3.1 and 1 µl 10 U/µl *Bsl*I (New England Biolab, USA). The reaction was performed at 37°C for 3 hours. The products obtained after digest were visualized by electrophoresis on an agarose gel.

### Chlorophyll fluorescence measurements

To measure maximum photosystem II (PSII) efficiency, the FlourCam7 version 1.2.5.3 system (PSI, Czech Republic) was used. Firstly, seedlings were dark-adapted for 15 min to determine the minimum chlorophyll fluorescence yield (F_O_ value). Then, plants were exposed to a saturating flash light, leading to the maximum fluorescence emission (F_M_ value). The maximum PSII value was calculated as the ratio F_V_/F_M_, where F_V_ is the variable fluorescence (F_V_ = F_M_ – F_O_).

### Determination of pigments

To determine pigment contents, 20-30 mg of seedling shoots were collected and finely ground in liquid nitrogen. Then, plant materials were homogenized with 200 µl of Acetone/Tris buffer (80% acetone in 10 mM Tris-HCl pH7.8) by vigorously mixing. The mixture was incubated for 10 min in a cold ultrasonic bath before centrifugation at 13000 rpm for 10 min at 4°C. Next, the supernatant was transferred into a new centrifuge tube. The absorption values of the supernatant were measured using a NanoPhotometer® NP80 (Implen GmbH, Germany) at the following wavelengths: 470, 537, 647, 663, and 750 nm. Pigment concentration was calculated using the equations of Sims & Gamon (2002).

### RNA extraction and quantitative RT-PCR

For gene expression analysis, 5-day-old whole seedlings were ground in liquid nitrogen, and the total RNA was isolated using the RNeasy Plant Mini Kit (QIAGEN, Germany). 1 μg of total RNA was used to synthesize cDNA with the RevertAidTM H Minus First Strand cDNA Synthesis kit (Thermo Fisher Scientific, USA). The quantitative real-time PCR reaction was conducted using 1 µl cDNA (12.5 ng), 0.4 µl forward/reverse primer, 5 µl 2X SYBR green master mix (KAPA SYBR FAST Universal, KAPABIOSYSTEMS) and the CFX Connect Real-Time PCR Detection System (Bio-Rad, USA). The relative expression level was calculated using the method of Pfaffl (2001), in which three reference genes were used: *AtACTIN2, At5g46630,* and *AtPP2AA2*. The gene identifiers (IDs), gene names, gene-specific primers, and amplicon sizes are provided in Table S5.

### RNA sequencing

Total RNA was isolated using the RNeasy Plant Mini Kit (QIAGEN, Germany). Then, RNA quantity and quality were determined using the Agilent RNA 6000 Pico Kit with Bioanalyzer 2100 system (Agilent Technologies, USA). Samples with high purity and RNA Integrity Number (RIN) were sent to Novogene Co., Ltd (UK) for library preparation, RNA sequencing, and bioinformatics analysis according to their standard protocols. RNA-seq was done using an Illumina-based NovaSeq 6000 system with 150 bp paired-end read sequencing. Reads were checked with FastQC and cleaned up before mapping to the reference *A. thaliana* genome (TAIR10) by HISAT2 (Kim *et al*., 2019). The number of reads for each gene was counted using featureCounts of HISAT2. Then, the fpkm value (fragments per kilobase of transcript per million mapped reads) was calculated as the relative expression level of a gene in the sample, in which a gene is considered as expressed if the fpkm > 1. Differently expressed genes (DEGs) between the mutant and the wild type were determined using raw read counts by DESeq2 R package. Functional annotation of DEGs was performed using the gene ontology enrichment tool (Ashburner et al., 2000; Aleksander et al., 2023).

### Quantification of glucosinolates

3-5 mg of *A. thaliana* whole seedlings were ground in liquid nitrogen and subsequently mixed with 450 μl Methanol:Chloroform (2:1 volume ratio) extraction solution supplemented with 20 µg/ml sinigrin as an internal standard and 200 µl deionized water. Samples were incubated at room temperature for 60 min, followed by centrifugation at 10,000 x g for 20 min. The upper aqueous phase was used for GSL determination via Liquid Chromatography coupled with tandem Mass Spectrometry (LC-MS/MS). The separation of substrates was performed at 35°C with a Nucleoshell RP18 column (50 × 3mm, particle size 2.7 μm; Macherey-Nagel, Germany) using an Agilent 1290 High-Pressure Liquid Chromatography (HPLC) system. As eluents A and B, water and acetonitrile each containing 0.1% formic acid, were used respectively, with 0.5 ml/min flow rate. Initially, the separation of GSL was taking place with 2% of eluent B for the first 0.5min, which was increased linearly to 95% over the next 7min. After washing, the column remained at 95% eluent B for 3min. The starting conditions were restored within the next 0.5min and the column was re-equilibrated with 2% B for 2 min. The analytes were detected by Electrospray Ionization MS/MS (ESI-MS/MS) by an API 3200 triple-quadrupole LC-MS/MS system with an ESI Turbo Ion Spray interface and operated in the negative ion mode (AB Sciex, Germany). The ion source parameters were set as follows: 40psi curtain gas, -4000 V ion spray voltage, 650°C ion source temperature, 60psi nebulizing and drying gas. GSL-specific signals were acquired via Multiple Reaction Monitoring (MRM), with scan time of 15msec; Q1 and Q3 masses (Q1, Q3 resolution unit) and compound specific parameters for each analyte are described in Table S6. Peak areas were calculated automatically by IntelliQuant algorithm of the Analyst 1.6 software (AB Sciex, Darmstadt, Germany), with manual adjustment when necessary. GSL content in each sample was calculated in Microsoft Excel, after normalization to the internal standard and to fresh weight.Peak areas were calculated automatically by the IntelliQuant algorithm of the Analyst 1.6 software (AB Sciex, Germany), with manual adjustment when necessary. GSL content in each sample was calculated and normalized to the internal standard and the fresh weight.

### Determination of myrosinase activity by spectrophometric method

To determine the activity of myrosinase, around 100 mg of fresh plant materials were frozen in liquid nitrogen and quickly ground into powder. The ground sample was homogenized in 1 ml extraction buffer containing 10 mM of postassium phosphate (pH 6.5), 1 mM EDTA, 3 mM dithiothreitol (DTT), 5% glycerol, and protease inhibitors. After centrifuging at 12000 x g for 15 min at 4°C, the supernatant was filtered by passing through syring filters Chromafil xtra MV-45/25 0.45 µm and aquenous solution was collected into a new tube. The extraction was repeat for another two times and all combined. Then, the extracted solution was concentrated 10 times using Vivaspin® Turbo 4 PES 10 kDa MWCO (Sartorius, UK). Total protein concentration was determined using the ProtaQuant^TM^ Assay (Serva, Germany).

Myrosinase assay was quantified as the rate of sinigrin degradation by the concentrated protein extract. The reaction buffer contained 33.3 mM postassium phosphate (pH 6.5) and 0.1 mM sinigrin. After equilibrating the reaction buffer at 37°C for 15 min, the reaction started by adding 50 µg of extracted protein into 1 ml of reaction buffer. The decrease in absorbance at 227 nm of the reaction mixture was recorded at 37°C for 30 min. Myrosinase activity was calculated using the formular (A_0_ - A_t_)/(*E*\**t*), where A_0_ and A_t_ are the initial absorbance and the absorbance after time *t*, respectively; *t* is the reaction time (min); *E* is the molar extinction coefficient, 7500 for sinigrin (Piekarska *et al*., 2013). The final myrosinase activity is given as nmol of sinigrin degraded by 1 mg of concentrated protein extract per minute.

### Phylogenetic analysis

Putative WHIRLY proteins were identified using BLASTP (Altschul *et al*., 1990) and HMMER v.3.3.2 (Finn *et al*., 2011) based on the conserved Whirly domain of AtWHIRLY1 (Pfam: PF08536). First, the BLASTP search with a threshold E-value of 0.01 was used for initial identification of the potential WHIRLY protein available in database. Then the raw HMM profile was downloaded by searching the WHIRLY family in the Pfam database (Mistry *et al*., 2021), and the WHIRLY proteins were extracted from the protein database by hmmsearch with an E-value threshold of 1E-5. Finally, The Pfam was used to verify the presence of a Whirly topology (β-β-β-β-α-linker-β-β-β-β-α) in their protein structures. Redundant protein sequences, misannotated, and poor-quality sequences were filtered out. Multiple sequence alignment of putative WHIRLY domains was performed by using the ClustalW with default parameters (ref). The alignment was trimmed to the Whirly domain region and then used to build a phylogeny tree of the WHIRLY family. The rooted tree was constructed by using MEGA version X (Kumar *et al*., 2018) with the neighbor-joining (NJ) method and bootstrap analysis with 1000 replicates. Putative sequences from bacterial and archea were used as outgroups.

### Statistical analysis

Physiological measurements, qRT-PCR, and GSL quantification were performed with three to four independent biological replicates. The mean of biological replicates was used to calculate the standard deviations and standard errors. A two-tail paired Student’s *t*-test was performed to estimate the statistically significant differences between lines using Microsoft Excel. Asterisks indicate statistical significances with * for p-value < 0.05, ** for p-value < 0.01, and *** for p-value <0.001.

## Data availability statement

All relevant data supporting the results of this article can be found within the manuscript and the supporting materials provided. Raw data from the RNA-seq samples have been submitted to the National Centre for Biotechnology Information (NCBI) Sequence Read Archive (SRA) database (# PRJNA1101217). The data will be publicly available and accessible at https://www.ncbi.nlm.nih.gov/sra/PRJNA1101217. For any further information regarding the RNA-seq data, please contact the corresponding author.

## Acknowledgement

We would like to thank Siska Herzklotz and Nancy Zimmermann for their excellent technical support. This research was funded by the Deutsche Forschungsgemeinschaft (DFG, German Reasearch Foundation) – 400681449/GRK2498.

## Author contributions

NTL and HK conceived and designed the experiments. KS and HG generated the mutants. GA and KK were screening the mutants. NTL performed phenotypic measurement, RNA-seq analysis, myrosinase assay, phylogenetic analysis, and data analysis. MP, ZJ, and AS aided with the setup, acquisition, and GSL quantification. NTL generated the illustrations. NTL, HK, and KK wrote the manuscript. All authors read and approved the manuscript.

