## Supplementary figures and images for "WHIRLY1 regulates aliphatic glucosinolate biosynthesis in early seedling development of Arabidopsis"

### Figure S

**Figure S1**

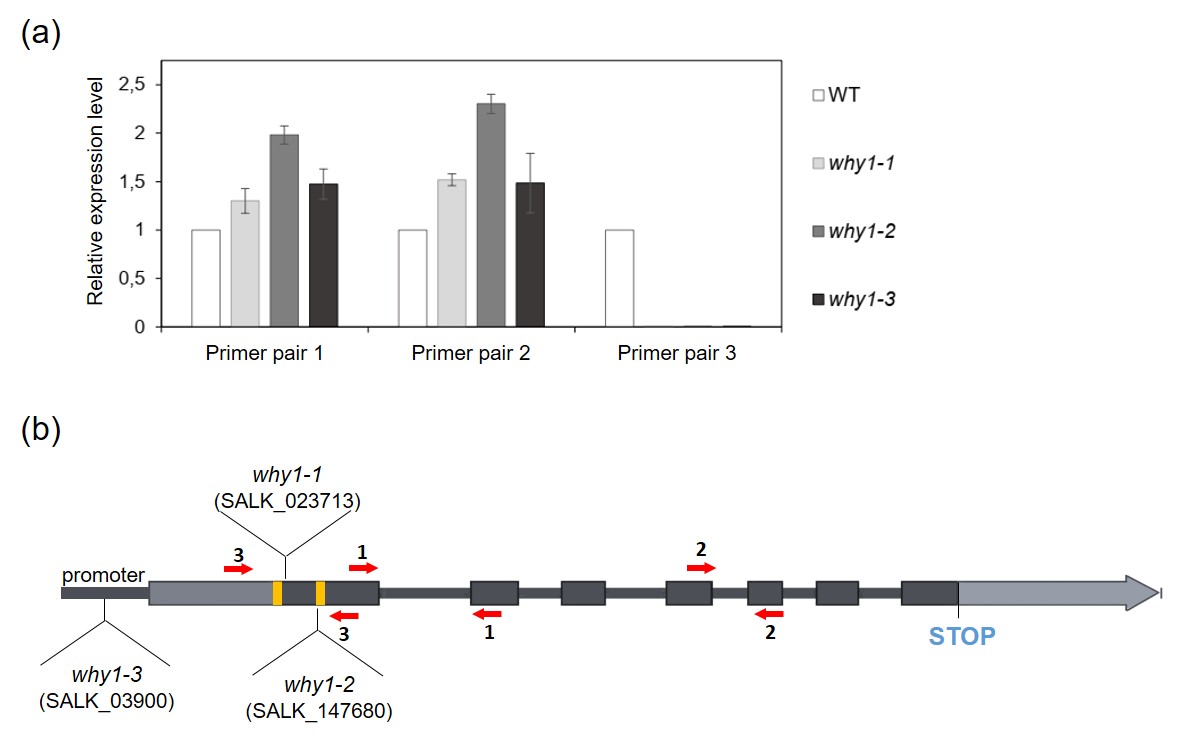

**Figure S2**

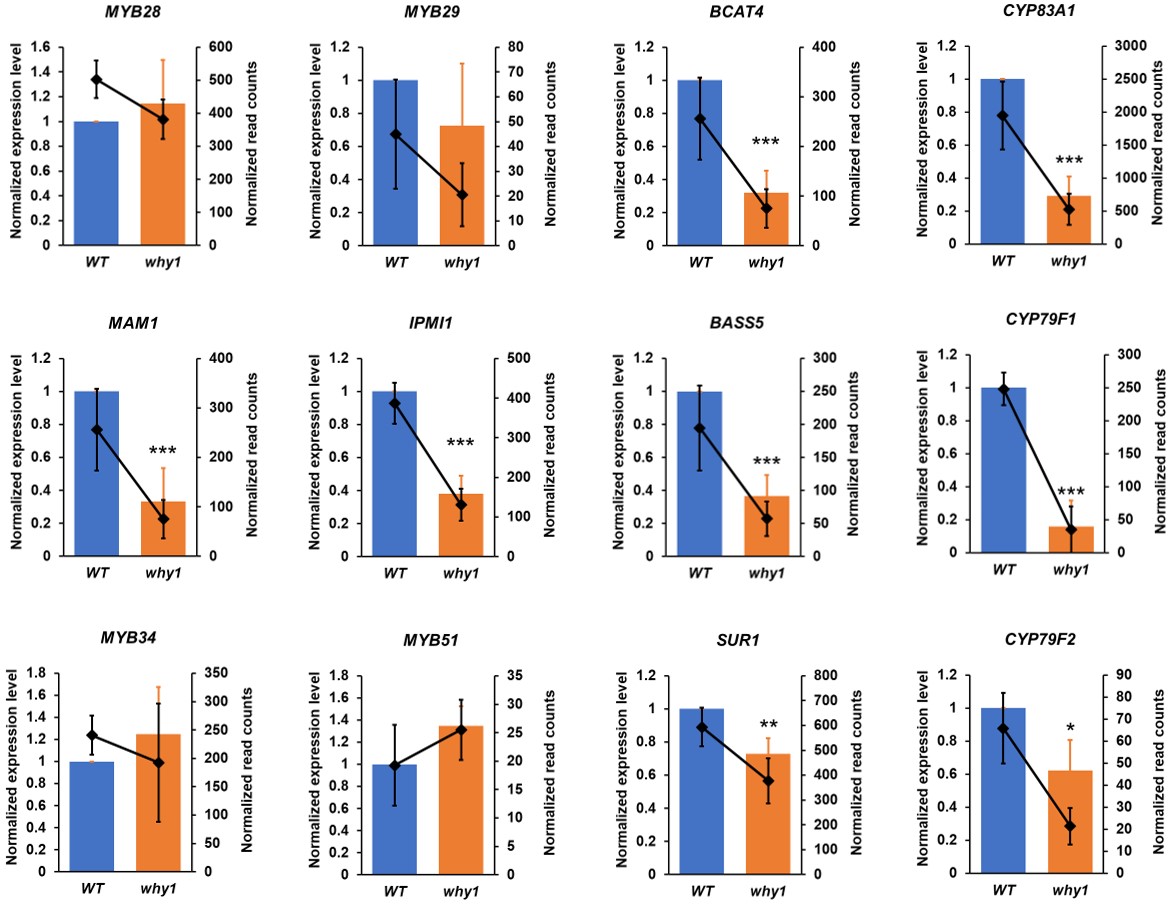

**Figure S3**

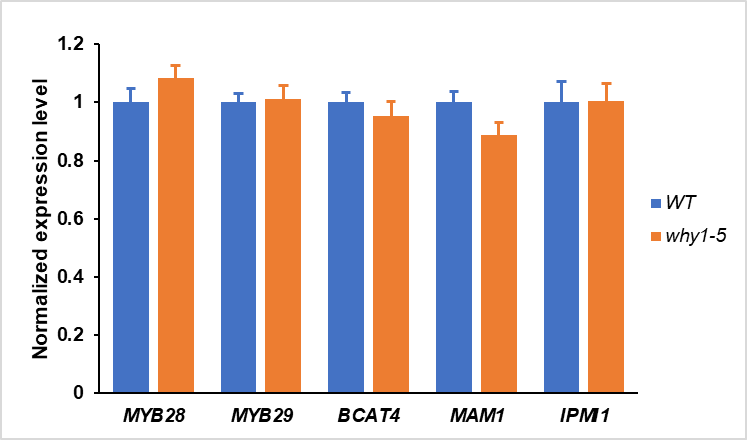

**Figure S4**

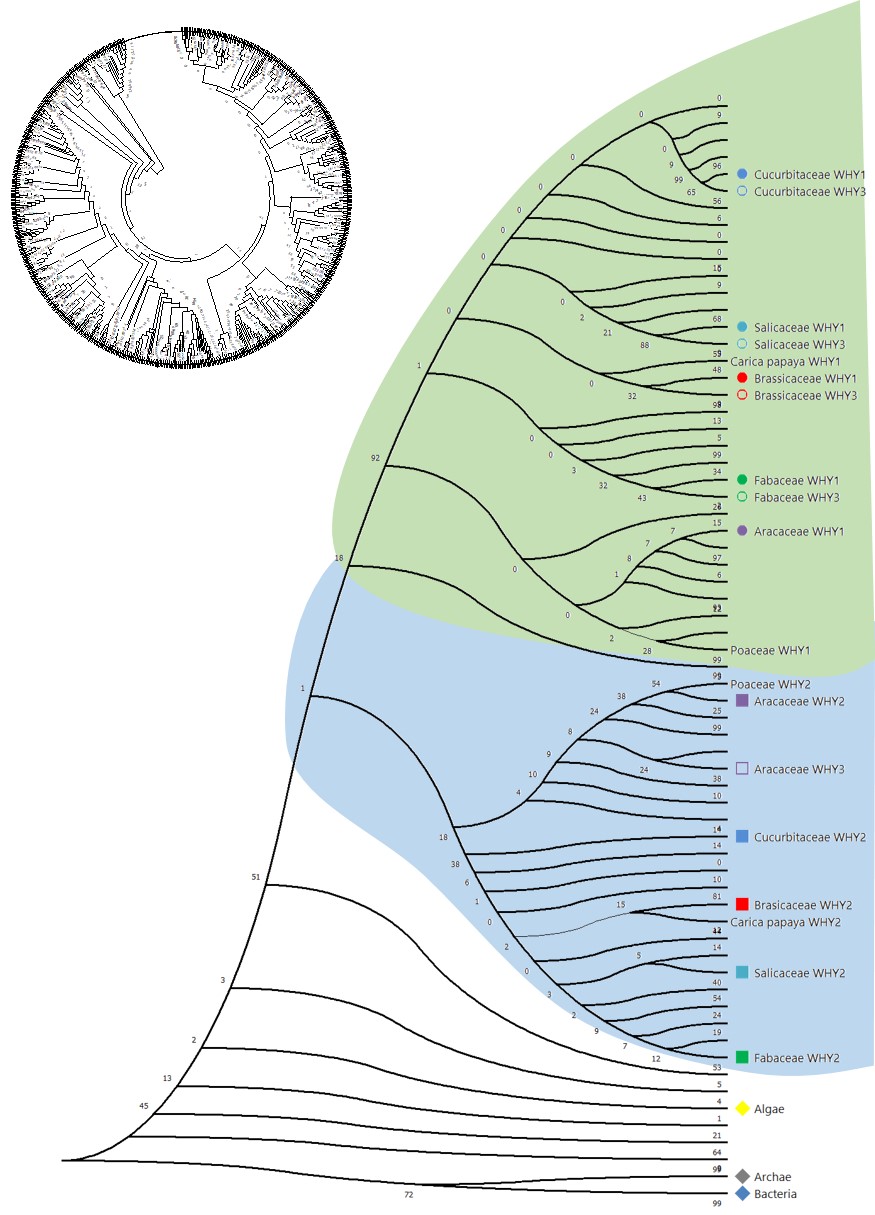

**Figure S5**

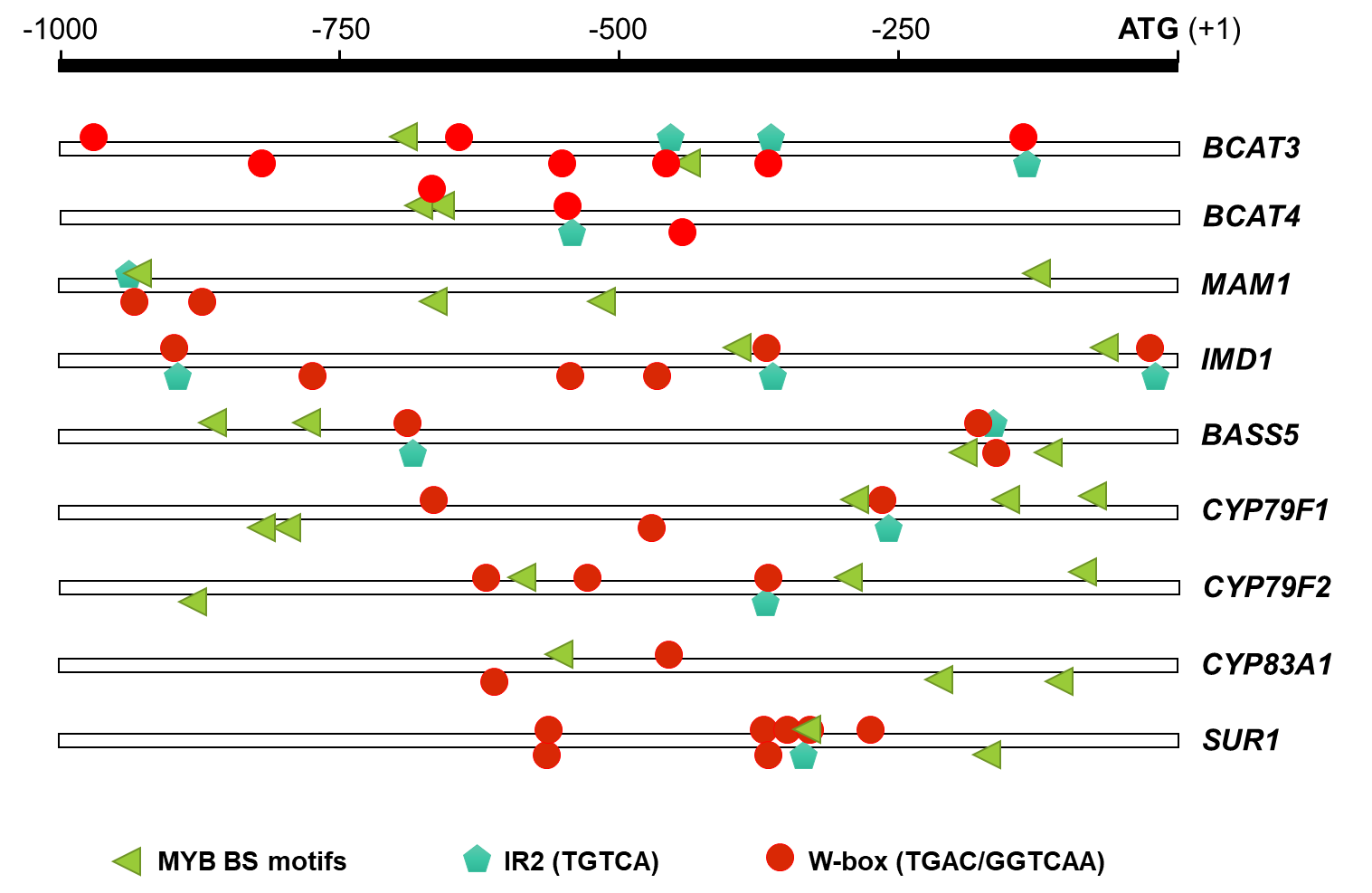
